# Temporal dynamics of BMP/Nodal ratio drive tissue-specific gastrulation morphogenesis

**DOI:** 10.1101/2024.02.06.579243

**Authors:** Alyssa A Emig, Megan Hansen, Sandra Grimm, Cristian Coarfa, Nathan D Lord, Margot Kossmann Williams

**Affiliations:** Center for Precision Environmental Health and Department of Molecular and Cellular Biology, Baylor College of Medicine, Houston, TX; Dan L Duncan Comprehensive Cancer Center, Baylor College of Medicine, Houston, TX; Department of Computational and Systems Biology, University of Pittsburgh, Pittsburgh, PA; Previous address: Department of Developmental Biology, Washington University School of Medicine, St. Louis, MO

**Keywords:** Gastrulation, Morphogenesis, Convergent extension, Zebrafish, Nodal, BMP

## Abstract

Anteroposterior (AP) elongation of the vertebrate body plan is driven by convergence and extension (C&E) gastrulation movements in both the mesoderm and neuroectoderm, but how or whether molecular regulation of C&E differs between tissues remains an open question. Using a zebrafish explant model of AP axis extension, we show that C&E of the neuroectoderm and mesoderm can be uncoupled *ex vivo*, and that morphogenesis of individual tissues results from distinct morphogen signaling dynamics. Using precise temporal manipulation of BMP and Nodal signaling, we identify a critical developmental window during which high or low BMP/Nodal ratios induce neuroectoderm- or mesoderm-driven C&E, respectively. Increased BMP activity similarly enhances C&E specifically in the ectoderm of intact zebrafish gastrulae, highlighting the *in vivo* relevance of our findings. Together, these results demonstrate that temporal dynamics of BMP and Nodal morphogen signaling activate distinct morphogenetic programs governing C&E gastrulation movements within individual tissues.

**SUMMARY STATEMENT:** Using zebrafish embryo and explant models, we demonstrate that temporal dynamics of morphogen signaling ratios distinguish between tissue-specific morphogenetic programs during vertebrate body plan formation.

## INTRODUCTION

During an embryo’s transition from a single cell to a complete organism, the nascent body plan first emerges through a series of highly conserved cell movements during gastrulation. It is here that the hallmark of the vertebrate body plan, the extended anteroposterior (AP) axis, is established via convergence & extension (C&E) movements that simultaneously narrow and elongate embryonic tissues (*1–4*). C&E of both the neuroectoderm (NE) and underlying mesoderm contribute to embryonic axis extension (*5, 6*), with each tissue exhibiting characteristic mediolateral (ML) cell elongation, alignment, and intercalations (*7–14*). While many aspects of C&E movements are similar between these tissues, they exhibit tissue-specific differences in the underlying cell behaviors. For example, distinct cellular mechanisms driving C&E of the epithelial NE vs. mesenchymal mesoderm in mice (*10, 11*), or the notochord boundary as a source of polarity cues in anamniote mesoderm (*8, 9, 15–18*). Spatiotemporal regulation of cell signaling is essential for proper morphogenesis, and both vertical and planar interactions between the mesoderm and NE influence C&E cell behaviors in each (*19–25*). This raises the question of whether a common morphogenetic program simultaneously drives C&E of both tissues, or whether each follows a distinct set of molecular instructions.

Answering such questions requires the ability to evaluate NE C&E independent of these movements in the mesoderm and vice versa, necessitating physical and chemical isolation of embryonic tissues. *Xenopus* dorsal marginal zone (DMZ) explants (*26*) and zebrafish whole-blastoderm explants (aka “pescoids”) (*27–29*) both undergo C&E, but contain endogenous signals present within the whole embryo. While C&E cell behaviors have been characterized in NE and mesoderm isolated from *Xenopus* DMZ (*7–9*), such tissues are likely not fully independent due to the exchange of planar signals prior to explantation (*21*). To overcome some of these limitations, we generate explants from only the animal blastomeres of zebrafish embryos to produce relatively naïve clusters of embryonic cells (*30–34*), similar to *Xenopus* animal cap explants (*35, 36*). In response to exogenous activation of the transforming growth factor β (TGF-β) family Nodal pathway, these explants specify all three germ layers and recapitulate the polarized cell behaviors and C&E movements observed in zebrafish gastrulae (*32–34*). We previously showed that extending explants injected with *ndr2* mRNA (encoding the Nodal ligand Cyc) exhibit mesoderm marker expression, but this domain was typically restricted to one end of each explant (*32*). The elongated portion instead expressed markers of NE (*32*), implying that their extension results largely from C&E cell behaviors within the NE, thus enabling examination of NE morphogenesis uncoupled from underlying mesodermal tissues.

The ability of the Nodal pathway to induce both mesoderm specification and axis extension in explants is consistent with its role *in vivo*, where Nodal is necessary for both (*37–40*). For example, mouse embryos mutant for Nodal signaling components fail to gastrulate entirely (*41*). Zebrafish embryos lacking all Nodal function - through loss of the coreceptor Tdgf1/Cripto (MZ*oep-/-*) (*40*), ligands (*sqt*-/-*cyc*-/-) (*38*), or downstream effector Smad2 (MZ*smad2*-/-) (*42*) – similarly lack all endoderm and most mesoderm and undergo abnormal gastrulation movements resulting in a severely shortened AP axis. Axis extension of MZ*oep*-/- embryos can be partially rescued by the restoration of axial mesoderm (*23*), demonstrating that Nodal signaling within the mesoderm alone is sufficient to drive extension of the embryonic axis, including the overlying NE. Indeed, mesoderm is known to influence morphogenesis of the overlying NE in zebrafish and *Xenopus* via direct coupling through the ECM (*24*), friction forces (*22*), and signaling molecules (*19, 43*). Together, these findings led to the assumption that the role of Nodal signaling in AP axis extension is largely secondary to its role in mesoderm specification (*23, 25*). This was reinforced by findings in *Xenopus* explants, in which mesoderm formation and C&E were so closely associated that one was proposed as a proxy for the other (*35*). However, the functions of Nodal in cell fate specification and morphogenesis can be distinguished. Non-Nodal signals can induce mesoderm specification within *Xenopus* explants without inducing C&E (*44–46*) and conversely, some signals promote explant C&E in the absence of mesoderm (*47, 48*). In *Xenopus* gastrulae, a subset of Nodal ligands promote C&E of the mesoderm independent of its specification (*49*), demonstrating the independence of these two processes. We further found that Nodal promotes ML cell polarity and C&E cell behaviors cell-autonomously within the NE of zebrafish embryos (*32*). These findings highlight a role for Nodal signaling in morphogenesis independent of cell fate specification, raising questions as to how Nodal promotes C&E of the NE and how or whether this molecular program differs from Nodal-induced mesoderm C&E.

The classic morphogen threshold model posits that outcomes like cell fate specification depend on morphogen signaling levels alone. However, countless studies reveal a more complex reality in which fate specification can rely on the cumulative dose (or time integral) of morphogen exposure (*50–53*), rate of change in morphogen concentration (*54, 55*), upstream signal activation dynamics (*56*), downstream kinetics of target gene expression (*42*), spatial fold change (*57–59*), competence windows (*60*), interactions between signaling pathways (*56, 61–64*), and others. Experimental manipulation of the timing, and thereby cumulative dose, of Nodal signaling disrupts mesendoderm patterning (*50, 53, 65*) and C&E (*32*). These dynamics are further complicated by interactions of Nodal with Bone Morphogenetic Protein (BMP) signaling, a fellow TGF-β pathway that induces ventral embryonic cell types (*66–69*) and inhibits C&E in zebrafish and *Xenopus* (*70–72*). These two signaling pathways interact at the level of both tissues and individual nuclei, with each affecting transcriptional responses of the other (*56*). For example, the BMP signaling gradient is in part restricted by the secreted BMP antagonist Chordin (*chrd*), a Nodal target gene (*29, 73*). Indeed, the ratio of these 2 signaling pathways governs activity of the gastrula organizer (*30, 74–78*) and is even sufficient to recapitulate an entire embryonic axis from naïve zebrafish explants (*30*). Together, these studies demonstrate that the ratio of BMP to Nodal signaling in both space and time during early embryonic development is essential for proper tissue specification and morphogenesis.

Here, we demonstrate that the temporal dynamics of BMP and Nodal signaling (and the ratio of the two) work in tandem to promote tissue-specific C&E morphogenesis during gastrulation. We find that naïve zebrafish explants can be induced to undergo either NE- or mesoderm-driven C&E by activation with a Nodal ligand (*ndr2*) or constitutively active Nodal receptor (CA-*acvr1b\**), respectively. Although these two molecules activate the same downstream effectors, their temporal dynamics differ substantially, with *CA-acvr1b\** inducing a robust Nodal response earlier than *ndr2*. Through direct optogenetic manipulation of signaling onset, we identify temporal windows during which Nodal signaling drives either mesoderm- or NE-driven C&E in explants. This response can be further modified by BMP signaling, with high BMP/Nodal ratios promoting NE C&E at the expense of mesoderm extension *ex vivo*. Finally, we demonstrate that increasing BMP/Nodal ratios during this critical window promotes C&E behaviors cell-autonomously within the ectoderm of intact embryos. Together, these data support a model by which BMP/Nodal ratios act during discrete temporal windows to promote distinct, tissue-specific C&E programs during gastrulation.

## RESULTS

### Different methods of Nodal signaling activation induce distinct, tissue-specific modes of C&E *ex vivo*

We previously showed that animal pole explants generated from zebrafish embryos injected with 10 pg *ndr2* mRNA (encoding the Nodal ligand Cyc) underwent robust C&E morphogenesis driven largely by the NE (*32*). To further investigate this NE-driven mode of C&E, we generated *ndr2* explants from *lhx1a:EGFP* transgenic embryos, in which the axial and lateral/intermediate mesoderm are labeled with EGFP (*79*). We further compared these to *lhx1a:EGFP* explants injected with 0.5 pg mRNA encoding a constitutively active Nodal receptor (*CA-acvr1b\**, also known as *taram-A-D* (*80*)) (Figure 1A). As expected, control explants from uninjected embryos exhibited no extension or mesoderm formation (as indicated by lack of EGFP expression) when intact sibling embryos reached the 4-somite stage (Figure 1B). Consistent with our previous study, *ndr2* explants exhibited robust extension and contained EGFP+ mesoderm domains that were restricted to one end of the explant, suggesting NE-driven extension (Figure 1C). *CA-acvr1b\** also induced EGFP+ mesoderm formation and robust extension, but in stark contrast to *ndr2* explants, the mesoderm domains of *CA-acvr1b\** explants were elongated along the axis of extension, indicating mesoderm-driven C&E (Figure 1E). When injected with a 10-fold higher dose (100 pg) of *ndr2*, many of explants were converted almost entirely to EGFP+ mesoderm, but few had extended by the equivalent of 4-somite stage (Figure 1D). This indicates that the observed NE- and mesoderm-driven modes of extension were not simply determined by Nodal activity levels but rather by the method of Nodal activation. Whole mount in situ hybridization (WISH) for mesoderm (*tbxta, noto*, and *tbx16*) and NE (*sox2, otx2*) marker genes further corroborated these findings (Figure 1G-K). Length measurements of *tbxta* expressing mesoderm domain as a proportion of the entire explant length indicate a significant shift toward mesoderm extension in *CA-acvr1b*\* compared with *ndr2* explants (Figure 1F). Notably, expression of the NE marker *sox2* differed between a striped pattern in *ndr2* explants, and diffuse staining in *CA-acvr1b\** explants (Figure 1J). Together, these results demonstrate that zebrafish blastoderm explants can be induced to undergo either mesoderm- or NE-driven axis extension, enabling examination of distinct C&E programs within morphogenetically uncoupled tissues.

**Figure 1:**
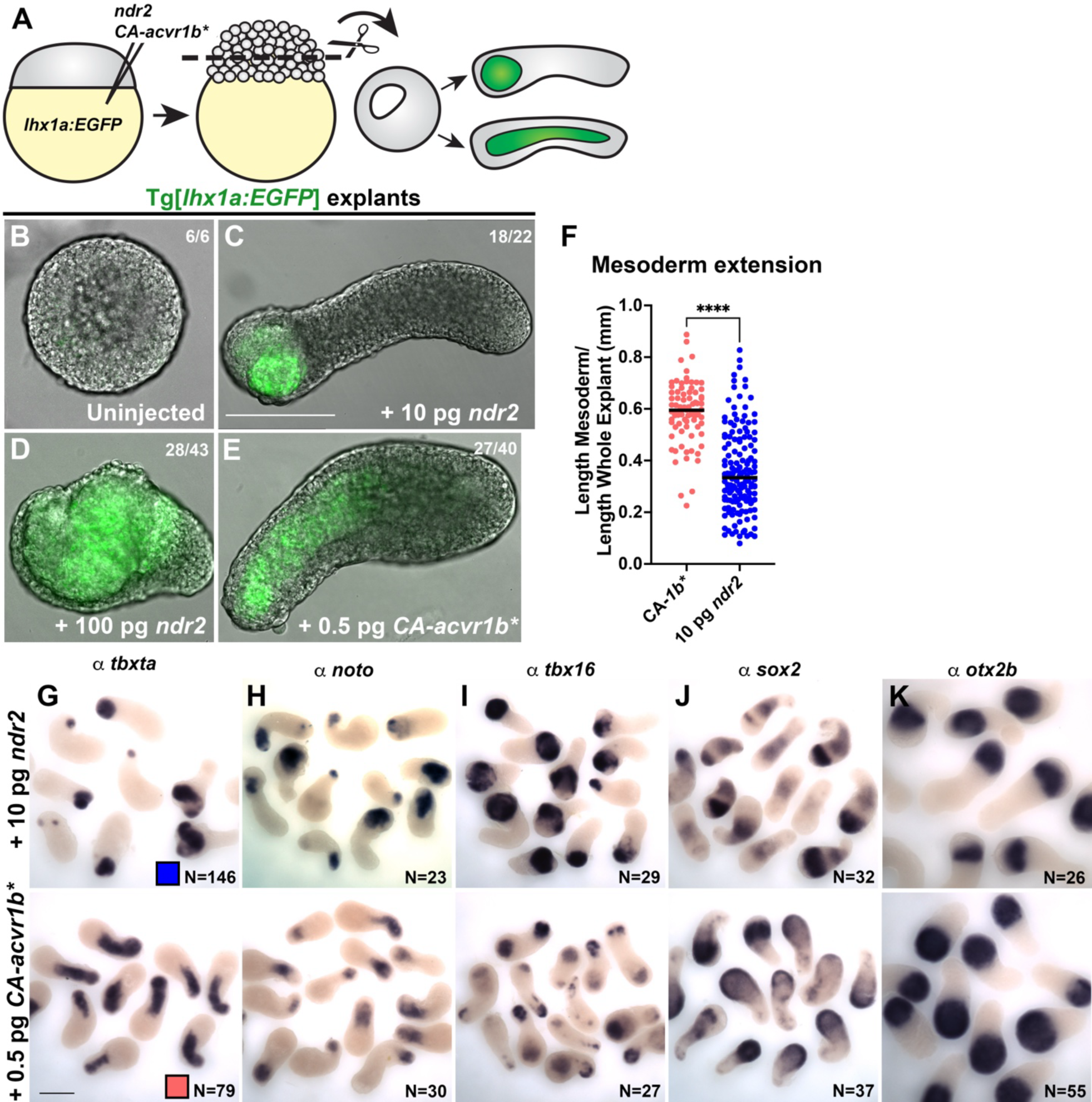
Different methods of activating Nodal signaling promote tissue-specific C&E in zebrafish explants. **A)** Diagram of Nodal injections and explantation of zebrafish embryos. **B-E**) Representative images of live Tg(*79*) explants, in which mesoderm expresses EGFP, of the conditions indicated at the equivalent of 4-somite stage. Fractions indicate the number of explants with the pictured phenotype over the number of explants examined. **F**) Mesoderm extension within explants as measured by the length of the *tbxta* mesoderm domain (shown in (G)) divided by the whole explant length. Each dot represents this ratio for an individual explant. Only explants with an overall length/width ratio of ≥1.4 are considered “extended” and thus, included in this analysis. Black bars are median values, ****p=<0.0001, Mann-Whitney test. The number of explants per condition is indicated in the corresponding image in (G). **G-K**) Representative images of WISH for the mesoderm markers *tbxta*, *noto*, and *tbx16* (G-I) and the neuroectoderm markers *sox2* and *otx2b* (J-K) in explants of the conditions indicated at the equivalent of 2-4 somite stage. N = number of explants examined per probe and condition from 5 independent trials. Scale bars = 200 μm.

### Longitudinal explant profiling reveals distinct temporal dynamics of Nodal and BMP signaling between tissue-specific C&E modes

Because *ndr2* and *CA-acvr1b\** are expected to activate the same (Nodal) signaling cascade, we next sought to determine the molecular mechanisms that distinguish the distinct morphogenetic outcomes elicited by each molecule. To this end, we compared transcriptomic profiles of *ndr2*-, *CA-acvr1b\**-, and uninjected control explants by bulk RNA-sequencing at each of seven developmental stages between sphere (4 hours post fertilization (hpf)) and 2-somite (11 hpf) (Figure 2A, Supplemental Dataset 1). Nodal signaling activity is tightly regulated in time and space during early zebrafish development (*42, 50*), prompting us to examine expression levels of 36 validated direct Nodal target genes (*42*) over time. By comparing Nodal target expression at each stage relative to their expression at 4 hpf, we observed strikingly different temporal patterns of Nodal-dependent transcriptional activity between explant conditions (Figure 2C). Nodal target expression in *CA-acvr1b\** explants increased steadily from 4.7 to 8 hpf, after which it plateaued (Figure 2C). By contrast, Nodal target expression in *ndr2* explants remained low until it rapidly increased at 8 hpf, nearly 3 hours after a similar increase in *CA-acvr1b\** explants (Figure 2C). This was corroborated by immunofluorescent staining against phosphorylated (p)Smad2, which first detected robust Nodal activity in *CA-acvr1b\** and *ndr2* explants at 4 and 8 hpf, respectively (Figure 2D,E). This suggests that an earlier or later surge in Nodal signaling activity may contribute to mesoderm-driven and NE-driven C&E modes, respectively.

**Figure 2:**
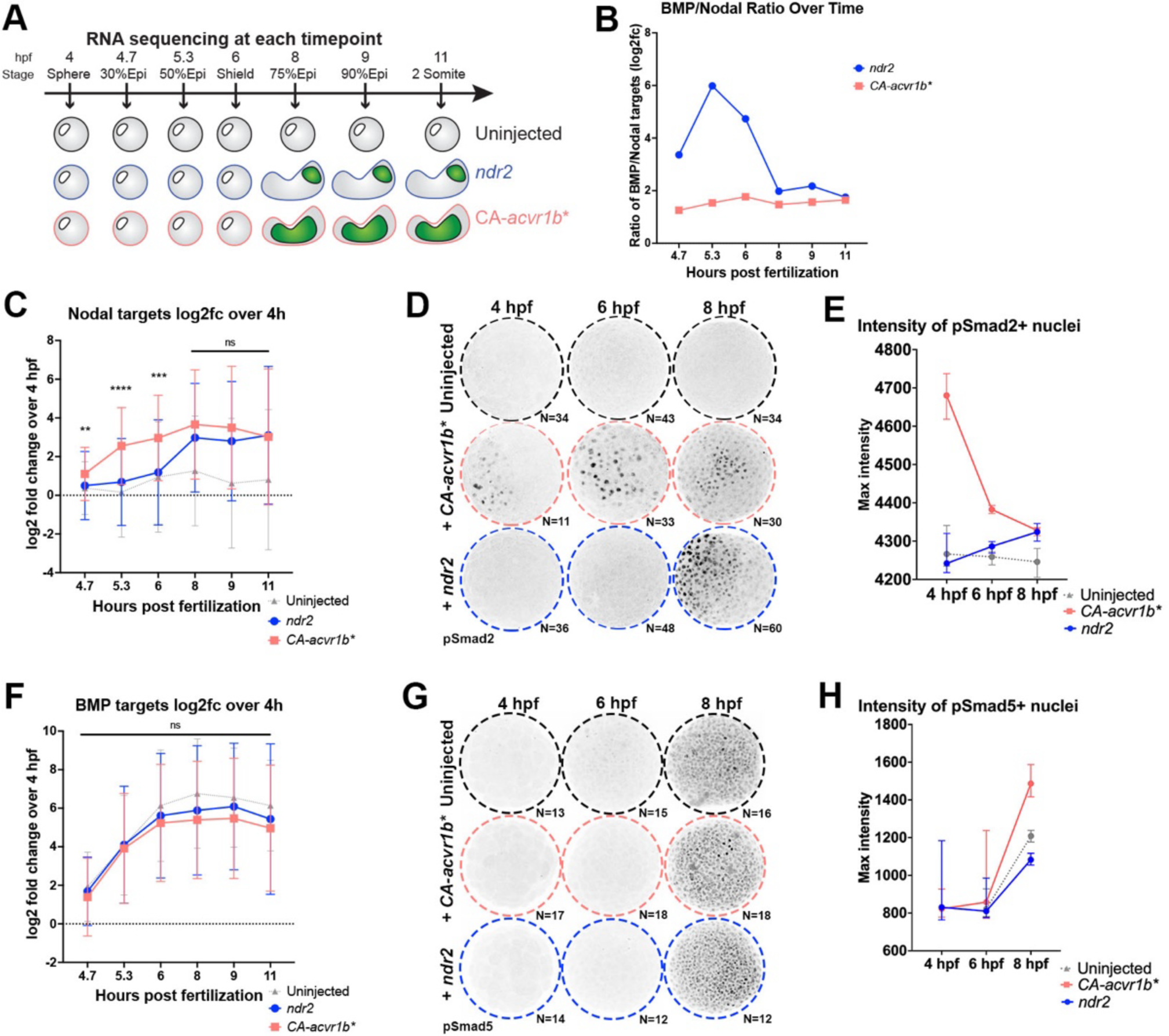
Nodal activity and BMP/Nodal ratios exhibit distinct temporal dynamics in *ndr2* and *CA-acvr1b\** explants. **A)** Overview of longitudinal transcriptional profiling in uninjected, *CA-acvr1b*,* and *ndr2* explants. **B**) Ratio of BMP- to Nodal-dependent transcriptional activity in *ndr2* and *CA-acvr1b*\* explants over time as estimated from the log2 fold change values of BMP and Nodal target gene expression in (C) and (F). **C**) Mean expression levels of 36 validated direct Nodal target genes in uninjected, *ndr2,* and *CA-acvr1b*\* explants at the time points indicated, represented as log2fold change over expression levels at 4 hpf. ****p=<0.0001, ***p=0.0002, **p=0.006, Mann-Whitney test of *ndr2* vs *CA-acvr1b*\*, error bars are standard deviation. **D**) Representative images of immunofluorescent pSmad2 staining in explants of the conditions and at the time points indicated. N= number of explants analyzed from 2 independent trials. **E**) Quantification of maximum staining intensity within pSmad2+ nuclei, shown in (D). Symbols are median values with 95% confidence interval. **F**) Mean expression levels of 11 validated direct BMP target genes in uninjected, *ndr2,* and *CA-acvr1b*\* explants at the time points indicated, represented as log2fold change over expression levels at 4 hpf. ns- Mann-Whitney test of *ndr2* vs *CA-acvr1b*\*, error bars are standard deviation. **G**) Representative images of immunofluorescent pSmad5 staining in explants of the conditions and at the time points indicated. N= number of explants analyzed from 1 trial. **H**) Quantification of maximum staining intensity within pSmad5+ nuclei, shown in (G). Symbols are median values with 95% confidence interval.

Although our relatively naïve blastoderm explants are isolated from endogenous Nodal, Wnt, and FGF signaling centers at the embryonic margin, they robustly express BMP signaling components (*30*). BMP activity antagonizes C&E movements during gastrulation (*70, 72*), and Nodal signaling can modulate BMP activity via the expression of the BMP antagonist *chordin* (*chrd*) (*29, 81–83*). Indeed, this interplay of Nodal and BMP signaling is sufficient to organize the entire zebrafish AP axis (*30, 56, 76, 77*). Reasoning that BMP signaling could modify the morphogenetic response of our explants, we next examined temporal expression patterns of validated direct BMP target genes (*61*) in our longitudinal RNA-seq dataset (Figure 2F). We found no significant difference in BMP target expression between explant conditions at any time point (Figure 2F), consistent with immunofluorescent staining for pSmad5 (Figure 2G,H), a cytoplasmic effector of BMP signaling. However, because Nodal target expression was significantly lower in *ndr2* explants at 5.3 and 6 hpf, their ratio of BMP to Nodal activity was substantially higher than *CA-acvr1b\** explants at these stages. Indeed, while the estimated BMP/Nodal ratio of *CA-acvr1b\** explants remained relatively low and essentially flat across development, *ndr2* explants exhibited a dramatic peak in BMP/Nodal ratio between 4.7 and 8 hpf (Figure 2B). This raises the possibility that temporal Nodal signaling dynamics and their interplay with BMP activity distinguish between mesoderm- and NE-driven C&E.

### Time of Nodal signaling onset determines the mode of tissue-specific C&E

To test the hypothesis that the timing of Nodal signaling activity determines the mode of tissue-specific C&E exhibited by our explants, we employed a newly-designed pair of optogenetic Nodal receptors (McNamara et al, in preparation) (hereafter, opto-Nodal) to precisely manipulate the onset of Nodal activity. These opto-Nodal receptors were constructed by fusing the intracellular domains of the type I and type II Nodal receptors Acvr1b and Acvr2b to plant-derived protein domains that dimerize in response to blue light (Cry2 and CibN, respectively (*84*)) (Figure 3A). Blue illumination of embryos injected with opto-Nodal mRNA produced widespread Nodal activity as evidenced by ectopic pSmad2 staining at 50% epiboly stage (5.3 hpf) and characteristic dorsalized phenotypes at 12 and 30 hpf (Figure Supplement 1A-E), whereas control embryos kept in the dark were phenotypically normal. Both pSmad2 immuno-staining and *ex vivo* extension were also apparent in explants cut from opto-Nodal injected embryos and exposed to blue light, while those kept in the dark failed to extend (Figure Supplement 1A,B,C,F). Ectopic pSmad2 staining was first detectable in opto-Nodal injected embryos and explants 30 minutes after onset of blue illumination (Figure Supplement 1D-F).

**Figure 3:**
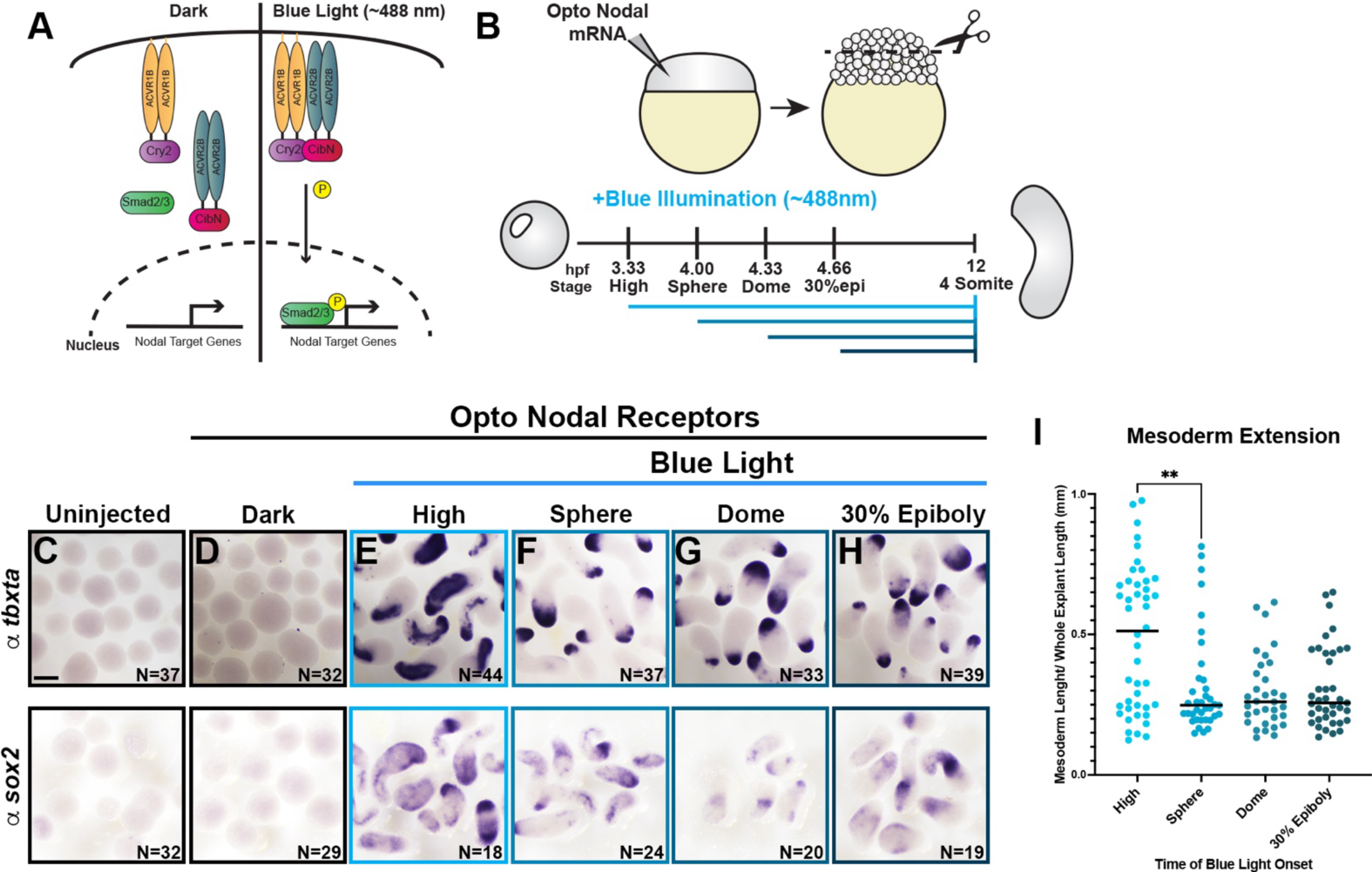
Timing of Nodal signaling onset is sufficient to determine tissue-specific C&E in explants. **A)** Diagram of optogenetic Nodal (opto-Nodal) receptors. **B**) Overview of opto-Nodal onset experiments in explants. **C-H**) Representative images of WISH staining for the mesoderm and NE markers *tbxta* (top) and *sox2* (bottom) in explants exposed to blue light at the stages indicated and analyzed at the equivalent of 4-somite stage. N = number of explants examined per probe and condition from 6 independent trials. Scale bar = 200 μm. **I**) Mesoderm extension within explants (shown in (E-H)) as described in Figure 1. Black bars are median values, **p=0.0053, Mann-Whitney test.

Having validated these tools in our *ex vivo* model, we next generated explants from opto-Nodal injected embryos and exposed them to blue light beginning at high (3.3 hpf), sphere (4 hpf), dome (4.3 hpf), or 30% epiboly (4.7 hpf) stages (Figure 3B). Illumination was maintained until intact siblings reached the 4-somite stage (12 hpf) (Figure 3B), at which point explants were fixed and evaluated for tissue-specific extension by WISH for mesoderm (*tbxta*) and NE (*sox2*) marker genes (Figure 3C-H). Many explants exposed to blue light at high stage contained extended mesoderm domains, as defined by the percentage of an explant’s length occupied by *tbxta* expression (Figure 3E, I). When we activated Nodal signaling at any of the subsequent stages, however, extension of mesoderm domains was significantly reduced compared with illumination at high stage (Figure 3F-I). Because of the approximately 30-minute delay between opto-Nodal illumination and pSmad2 detection, we estimate that the developmental window during which robust Nodal activity induces mesoderm-specific C&E closes around 4 hpf. This directly demonstrates that the timing of Nodal signaling onset is sufficient to determine the mode of tissue-specific C&E within our explant model, with earlier and later activation favoring mesoderm- and NE-driven extension, respectively.

**Figure Supplement 1:**
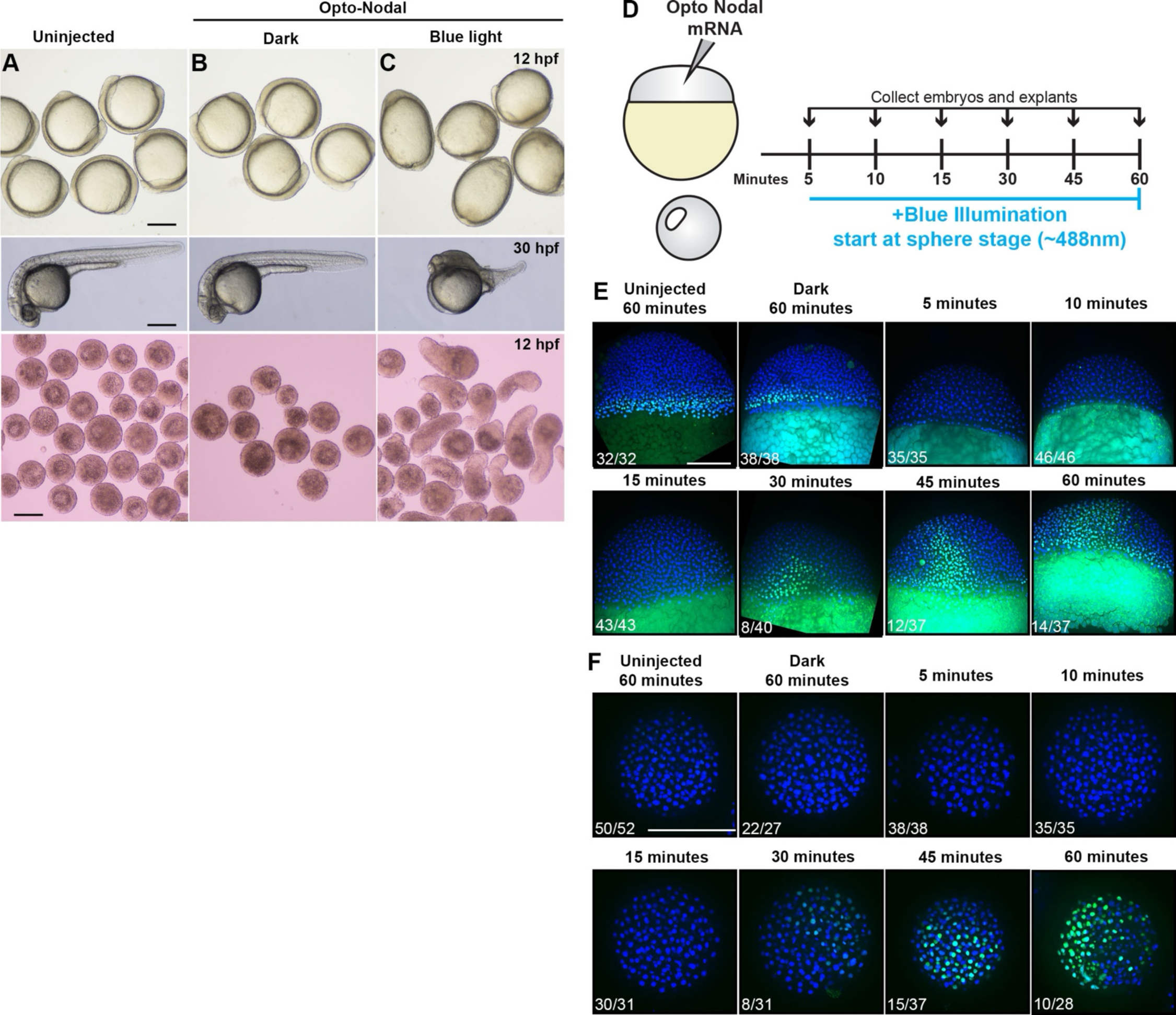
Validation of optogenetic Nodal receptors. **A-C**) Embryos at 12 hpf (4 somites) and 30 hpf and explants at 12 hpf that are uninjected (A), injected with 5 pg of mRNA encoding each Opto-Nodal receptor and kept in the dark (B), or placed under blue light at 3.5 hpf (high stage) (C). Note dorsalized phenotypes in (C). **D**) Overview of experiment to test delay between blue illumination and pSmad2 detection in embryos and explants injected with Opto-Nodal mRNA. Embryos and explants were placed under blue light starting at 4 hpf (sphere stage) and collected at different times after initial exposure. **E-F**) Immuno-staining for pSmad2 (green) in embryos (E) and explants (F) fixed at the indicated times after opto-Nodal activation. Blue counterstain is DAPI. Note the 30-minute delay between illumination and pSmad2 staining. Fractions indicate the number of embryos/explants with the pictured phenotype out of the total number examined. Scale bars = 200 μm.

### Increased BMP/Nodal signaling ratios switch explants from mesoderm- to NE-driven C&E

Our longitudinal transcriptional profiling revealed a large peak in the ratio of BMP/Nodal transcriptional activity in *ndr2* explants compared with *CA-acvr1b\** explants through 6 hpf (Figure 2B). To determine if this increased BMP/Nodal ratio is sufficient to promote NE-driven extension, we experimentally increased BMP activity within *CA-acvr1b\** explants (which otherwise exhibit mesoderm-driven extension) (Figure 4A). To this end, we co-injected mRNAs encoding a constitutively active BMPR1 receptor (*CA-BMPR1*) (*85*), the atypical BMP ligand anti-dorsalizing morphogenic protein (*admp*) (*86*), or multi-guide CRISPRs targeting the BMP antagonist *chrd* (Figure 4B, Figure Supplement 2A-D). Strikingly, all three manipulations caused *CA-acvr1b\** explants to switch their mode of extension from mesoderm- to NE-driven, exhibiting similar phenotypes to *ndr2* explants (Figure 4D-I). Control *tyrosinase* CRISPRs did not impact extension of mesoderm domains in CA-*acvr1b\** explants (Figure Supplement 2F-J).

**Figure 4:**
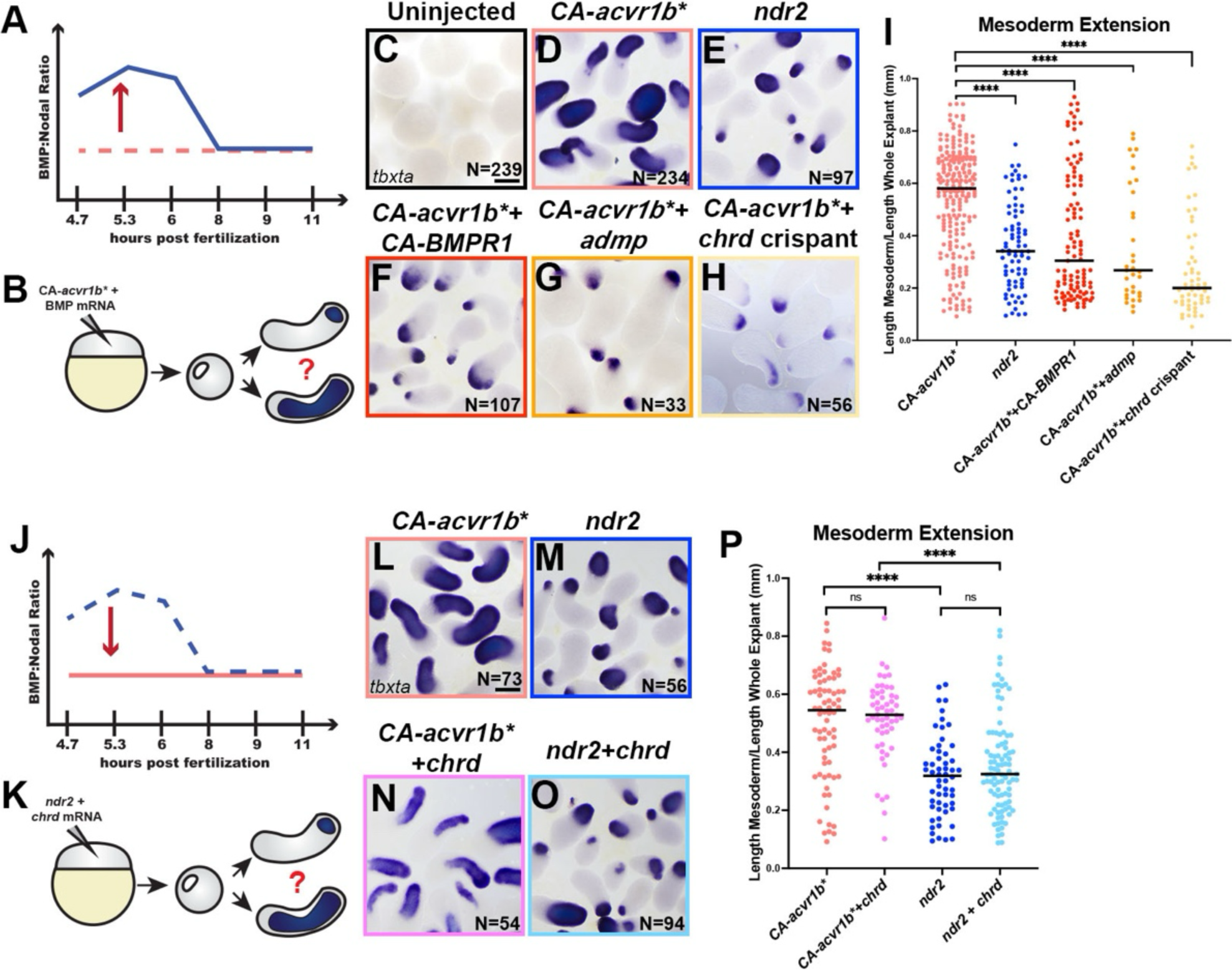
Increased BMP/Nodal ratios switch explants from mesoderm- to NE-driven C&E modes. **A-B**) Overview of experimental manipulation to make BMP/Nodal ratios in *CA-acvr1b\** explants (dotted line) resemble *ndr2* explants (solid line) (A) by injecting BMP modifiers *CA-BMPR1* mRNA, *admp* mRNA, and *chrd* CRISPRs (B). **C-H**) Representative images of WISH for *tbxta* in explants of the conditions indicated at the equivalent of 4-somite stage. N = number of explants examined from 3 independent trials. **I**) Mesoderm extension within explants (shown in (D-H)) as described in Figure 1. Black bars are median values, ****p= <0.0001 via Mann-Whitney test. **J-K**) Overview of experimental manipulation to make BMP/Nodal ratios in *ndr2* explants (dotted line) resemble *CA-acvr1b\** explants (solid line) (J) by injecting *chrd* mRNA (K). **L-O**) Representative images of WISH for *tbxta* in explants of the conditions indicated at the equivalent of 4-somite stage. N = number of explants examined from 3 independent trials. **P**) Mesoderm extension within explants (shown in (L-O)) as described in Figure 1. Black bars are median values, ****p= <0.0001, Mann-Whitney test. Scale bar = 200 mm.

In the converse experiment, we tested if decreasing the BMP/Nodal ratio in *ndr2* explants could similarly switch their mode of extension to mesoderm-driven (Figure 4J-K). We found that reducing BMP activity by co-injection with *chordin* mRNA (Figure Supplement 2D) had no effect on the mode of extension for either *ndr2* or *CA-acvr1b\** explants (Figure 4L-P), suggesting that reduced BMP/Nodal ratios are not sufficient to promote mesoderm-specific C&E without sufficiently high Nodal levels during early development. Together with our temporal manipulation of Nodal activity (Figure 3), these results indicate that high BMP/Nodal ratios promote NE C&E at the expense of mesoderm C&E, but that low BMP/Nodal ratios can only promote mesoderm C&E when Nodal signaling is high, and only during very early development.

**Figure Supplement 2:**
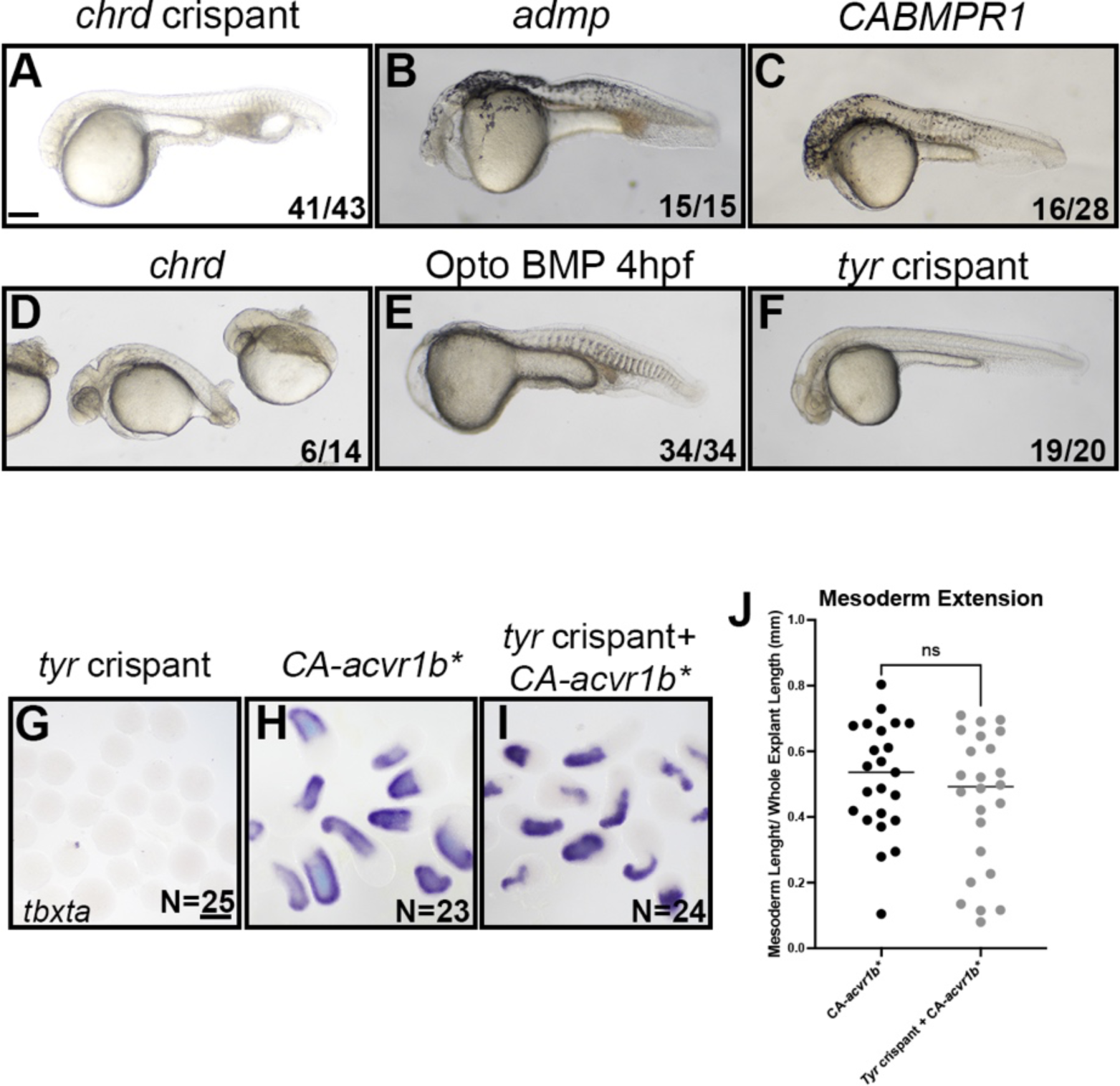
Validation of BMP modifiers. **A-F**) 24-30 hpf embryos injected with *chrd* gRNAs+Cas9 protein (A), *admp* mRNA (B), *CA-BMPR1* mRNA (C), *chrd* mRNA (D), *opto-BMP* mRNA and exposed to blue light at 4 hpf (sphere) (E), and *tyrosinase* gRNAs+Cas9 protein (F). Note the ventralized phenotypes in (A,B,C,E) and the dorsalized phenotype in (D). Fractions indicate the number of embryos/explants with the pictured phenotype out of the total number examined. **G-J**) Validation that control *tyrosinase* gRNA+Cas9 does not affect tissue-specific extension in explants. **J**) Mesoderm extension within explants WISH stained for *tbxta* (shown in (G-I)) as described in Figure 1. Black lines are median values and there was no significant difference in mesoderm extension between samples using a Mann-Whitney test. N= number of explants examined for each condition. Scale bars = 200 μm.

### BMP/Nodal ratio determines tissue-specific C&E mode during a critical temporal window

We showed above that different Nodal onset times elicited distinct morphogenetic responses in our explants (Figure 3). This raised the possibility of a similar temporal window during which BMP activity can modify tissue-specific C&E modes *ex vivo*. To test this, we utilized optogenetic BMP receptors (hereafter opto-BMP) (*61*) (Figure Supplement 2E) to activate BMP at increasingly later timepoints within *CA-acvr1b\** explants (Figure 5A,B), which otherwise exhibit low BMP/Nodal ratios and mesoderm-driven C&E. We found that opto-BMP activation at sphere and 50% epiboly stages (4 and 5 hpf) caused *CA-acvr1b\** explants to switch to predominately NE-driven extension (Figure 5C-F,J), as seen with other BMP signaling agonists (Figure 4). However, explants in which opto-BMP was activated at shield stage (6 hpf) or later exhibited no significant difference in extension of mesoderm domains when compared to *CA-acvr1b\** alone (Figure 5G-J). This demonstrates that increased BMP/Nodal ratios are only sufficient to switch explants from mesoderm- to NE-driven extension if applied prior to gastrulation stages, highlighting a critical developmental window during which BMP and Nodal establish tissue-specific morphogenetic programs.

**Figure 5:**
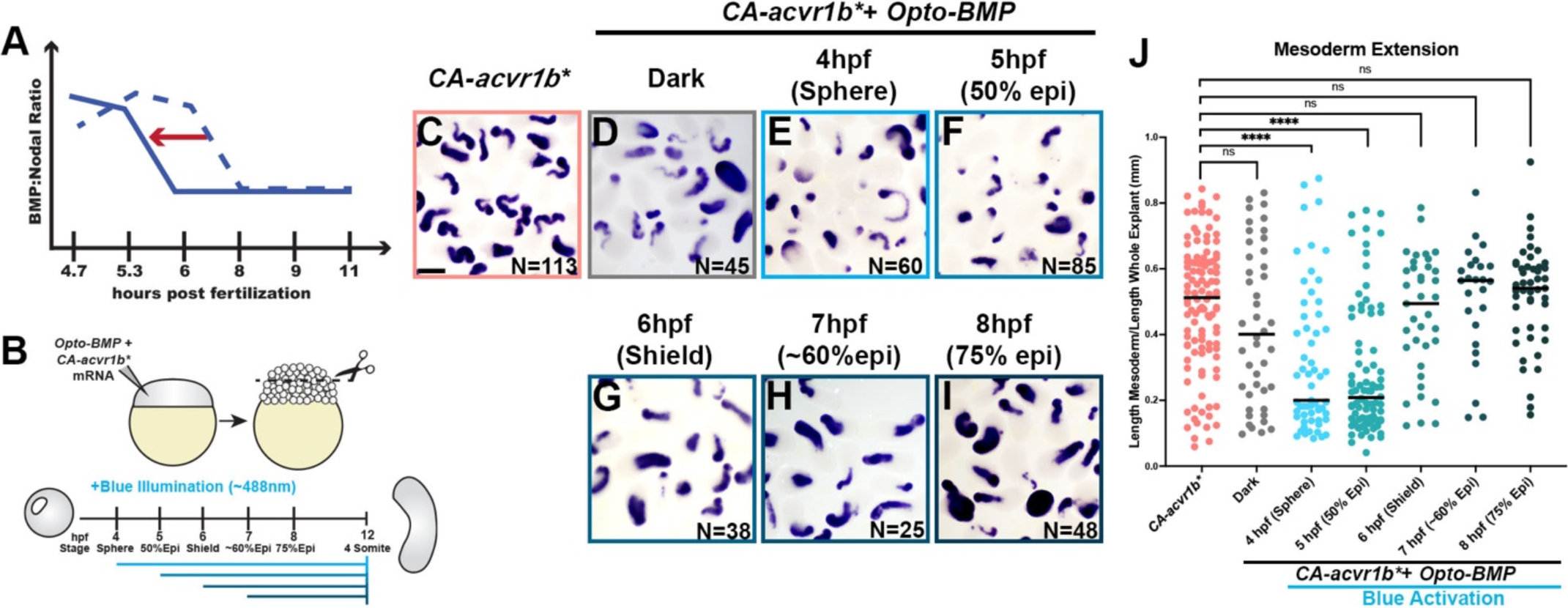
High BMP/Nodal ratios prior to gastrulation onset promote NE-driven C&E in explants. **A-B**) Overview of experimental manipulation to shift the temporal dynamics of BMP/Nodal ratios in *CA-acvr1b\** explants (A) by activating optogenetic BMP receptors using blue illumination at different time points (B). **C-I**) Representative images of WISH for *tbxta* in explants exposed to blue light at the time points indicated and analyzed at the equivalent of 4-somite stage. N = number of explants examined from 3 independent trials. Scale bar = 200 μm. **J**) Mesoderm extension within explants (shown in (C-I)) as described in Figure 1. Black bars are median values, ****p=<0.0001, Mann-Whitney test.

### Increased BMP/Nodal ratio promotes ectoderm C&E *in vivo*

To test whether temporal dynamics of BMP/Nodal signaling ratios similarly promote tissue-specific C&E within intact zebrafish gastrulae, we sought to increase BMP activity in a tissue-targeted fashion at different developmental stages. To this end, we transplanted the animal region, which is fated to form ectoderm, between opto-BMP-injected embryos and WT siblings at the 256-512 cell stage. (Figure 6A, Figure Supplement 3A). Control transplants between *lhx1a:EGFP* and WT embryos gastrulated normally (Figure Supplement 3B), with the donor animal region contributing to ectoderm and the host margin forming primarily mesoderm and some ectoderm (Figure Supplement 3C,D), validating our transplant approach for tissue targeting. In opto-BMP transplanted embryos, blue illumination was then used to increase BMP activity within primarily the mesoderm or ectoderm (Figure 6A). BMP activity was confirmed in *opto-BMP* + *H2B-RFP* donor animal poles exposed to blue light at 4 hpf (sphere stage), which exhibited robust pSmad5 staining that was absent from dark and WT donor controls (Figure Supplement 3E-H). When signaling was activated at 4 hpf (sphere stage), embryos expressing opto-BMP in the margin (mesoderm) appeared largely normal at gastrulation’s end (Figure 6B), whereas those with opto-BMP in the ectoderm became hyper-elongated (Figure 6C). This striking phenotype was quantified as an increase in the ratio of animal-vegetal length to dorsal-ventral width (Figure 6H), and resembled that of BMP signaling mutants in which C&E cell behaviors are expanded to the embryo’s ventral side (*66, 70*). Neither embryos kept in the dark, nor those in which opto-BMP was activated at 6 hpf (shield stage), exhibited a change in length/width ratios regardless of which tissue expressed opto-BMP (Figure 6 D-H). This is consistent with results from our explants, in which increased BMP promoted NE C&E but only before 6 hpf (Figure 5).

**Figure 6:**
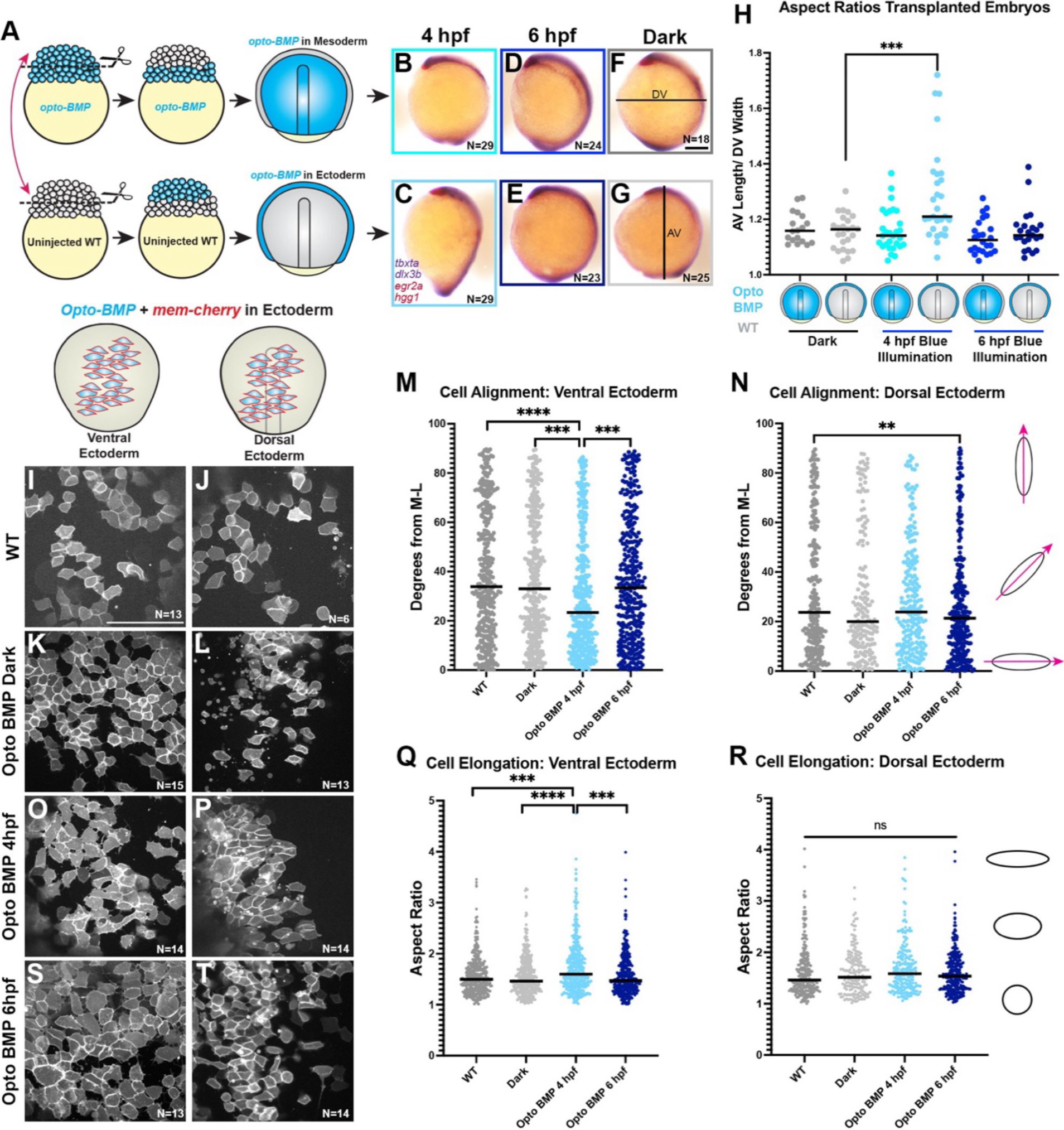
High BMP/Nodal ratios within ectoderm cells prior to gastrulation promotes C&E behaviors cell-autonomously. **A)** Diagram of animal pole transplants between *opto-BMP-* and un-injected embryos. **B-G**) Representative images of transplanted embryos with opto-BMP hosts (top) or donors (bottom) activated at 4 or 6 hpf (plus dark controls), then WISH stained for *tbxta*, *dlx3b* (purple), *eng2a*, and *hgg1* (red) at 2-somite stage (11 hpf). N = number of explants examined from 3 independent trials, scale bar = 200 μm. **H**) Ratio of animal-vegetal length (F) to dorsal-ventral width (G) of the embryos shown in (B-G). Each dot represents the aspect ratio of an individual embryo, black bars are median values, ***p=0.0005 Mann-Whitney test. **I-L, O-P, S-T**) Transplanted *opto-BMP/mem-Cherry*-expressing ectoderm in the ventral (I,K,O,S) and dorsal (J,L,P,T) sides of live embryos at tailbud stage (10 hpf). N = number of embryos from 3 independent trials, scale bar = 100 μm. **M-N, Q-R**) Cell alignment (M,N) and cell elongation (Q,R) of ventral (M,Q) and dorsal (N,R) ectoderm cells shown in (I-L, O-P, S-T). Cell alignment (M,N) is represented as degrees from the medio-lateral axis. Black bars are median values, ***p=0.0006, ****p=<0.0001, ***p=0.0008 (M), **p=0.0076 (N), Kolmogorov-Smirnov test. Cell elongation (Q,R) is represented as aspect (length/width) ratio of individual cells. Black bars are median values, ***p=0.0001, ****p=<0.0001, ***p=0.0004, Kolmogorov-Smirnov test. Each dot represents a single cell. Anterior is up in all images.

This result was surprising given that BMP signaling was found to reduce C&E behaviors cell-autonomously within the mesoderm (*70*). To determine whether BMP activity promotes *in vivo* ectoderm C&E cell-autonomously, we performed live confocal imaging of WT host embryos transplanted with animal poles expressing both opto-BMP and membrane Cherry. We then quantified ML alignment and elongation of labeled cells in both the dorsal and ventral embryonic regions at the end of gastrulation (10 hpf) after opto-BMP activation at 4 or 6 hpf (and in WT-WT and dark controls). As expected, we found that dorsal ectoderm cells were more mediolaterally aligned than ventral ectoderm cells in control embryos (Figure 6I-N), consistent with C&E and a lack thereof in these regions, respectively. Strikingly, opto-BMP activation at 4 hpf caused a significant increase in both ML cell alignment and elongation of ventral ectoderm cells, causing them to resemble dorsal ectoderm (Figure 6M-R). This effect was not seen when opto-BMP was activated at 6 hpf (Figure 6M,N,Q,R,S,T), consistent with length/width measurements of whole embryos (Figure 6H). This demonstrates that increased BMP signaling prior to gastrulation enhances C&E behaviors cell-autonomously within the ectoderm. Because BMP has the opposite effect on mesoderm cells (*70*), this finding supports our hypothesis that BMP/Nodal ratios elicit distinct morphogenetic effects in different tissues. Together with our explant experiments, these data are consistent with an early window during which BMP and Nodal levels determine tissue-specific morphogenetic outcomes, with high and low BMP/Nodal ratios promoting C&E of the ectoderm and mesoderm, respectively.

**Figure Supplement 3:**
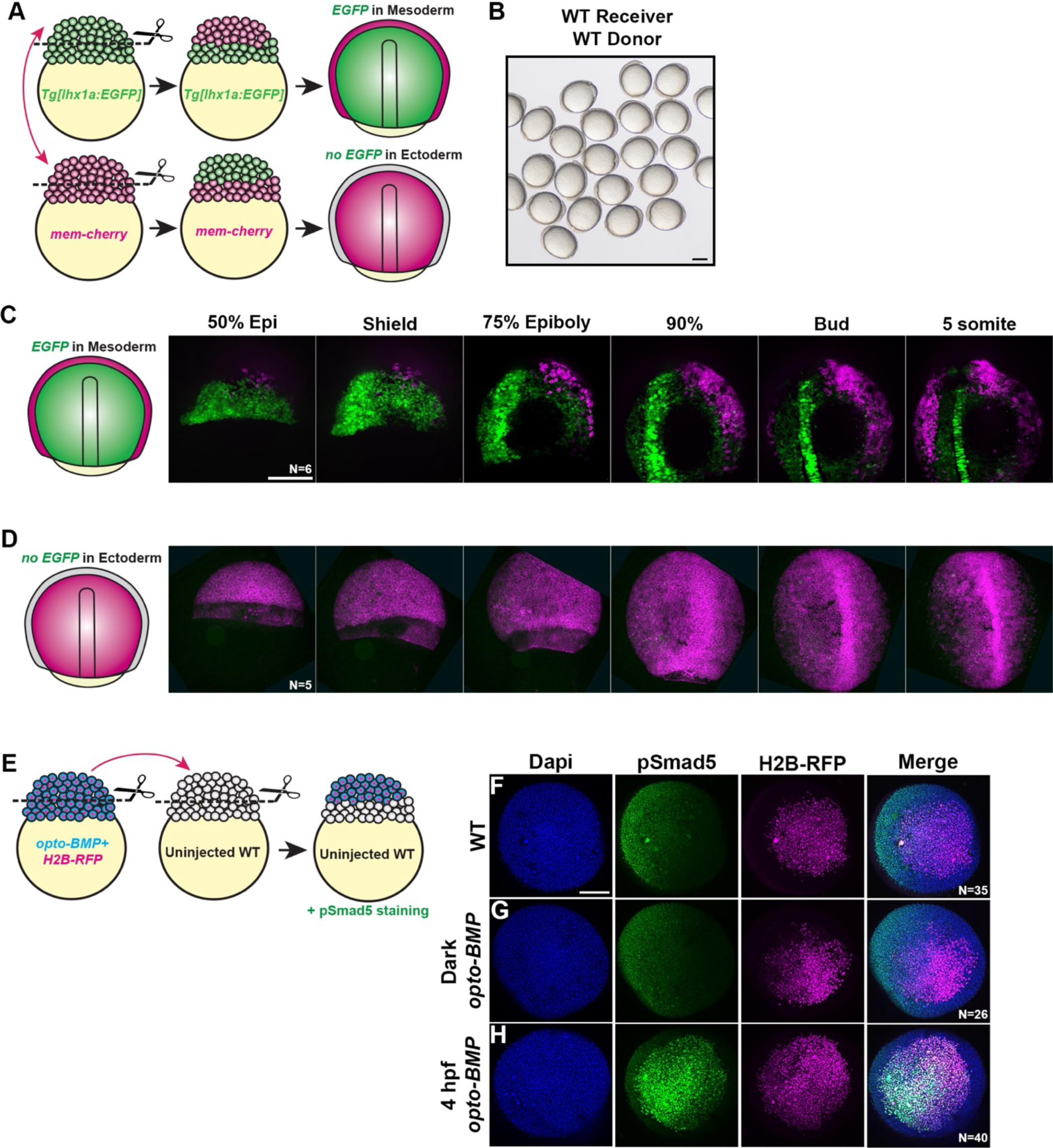
Characterization of animal pole transplants. **A)** Diagram of animal pole transplants between Tg(*79*) and *mem-cherry* expressing WT embryos. **B**) Transplanted WT to WT control embryos that were phenotypically normal at 11 hpf. **C-D**) Still images from representative 7-hour confocal time-lapse series of *mem-Cherry*+ animal poles transplanted onto Tg(*79*) embryos (C) and Tg(*79*) animal poles transplanted onto *mem-Cherry*+ embryos. Note that transplanted Tg(*79*) cells do not give rise to mesoderm. **E**) Diagram of animal pole transplants from an *opto-BMP* + *H2B-RFP* donor to an uninjected WT host embryo. After transplanting, embryos were collected at shield stage and immuno-stained for pSmad5 to detect BMP activity. **F-H**) Representative images of pSmad5 staining in shield-stage embryos transplanted with animal poles from *H2B-RFP* expressing WT (F) or *opto-BMP* donor embryos that were kept in the dark (G) or exposed to blue light at 4 hpf (sphere stage) (H). N= the number of embryos examined from each condition, scale bars = 200 μm.

## DISCUSSION

Coordinated morphogenesis of the mesoderm and overlying NE is essential for proper formation of the embryonic body plan during gastrulation. Both tissues undergo C&E to elongate the primary body axis, but it was unclear whether or how molecular regulation of morphogenesis differs between tissues. Here, we provide evidence for distinct C&E programs within the mesoderm and ectoderm that are determined by BMP/Nodal signaling ratios during a key developmental window. By activating Nodal signaling in otherwise naïve zebrafish embryonic explants, we demonstrated that morphogenesis of mesoderm and NE can be uncoupled *ex vivo*. Using this simplified system, we determined that Nodal signaling onset is sufficient to distinguish between tissue-specific extension modes, and that this effect can be modulated by BMP activity. We found that low BMP/Nodal ratios promote extension of the mesoderm, but only when Nodal levels are sufficiently high and only before 4 hpf. High BMP/Nodal, on the other hand, promotes NE C&E at the expense of mesoderm extension, and the window for this effect closes at gastrulation onset. Remarkably, these findings were recapitulated *in vivo*. BMP activation specifically within the ectoderm drove hyper-extension of intact gastrulae and caused a cell-autonomous expansion of C&E behaviors into normally non-extending regions of the embryo. As in explants, this effect of BMP was only observed before gastrulation onset, supporting a model in which the temporal dynamics of morphogen signaling activate tissue-specific C&E programs.

### Relevance of explants to intact embryos

Although our zebrafish explants contain all three germ layers and exhibit C&E cell behaviors, they do not faithfully recapitulate all aspects of gastrulation. For example, *ndr2* explants contain chordamesoderm, a tissue that undergoes robust C&E *in vivo*, but it fails to extend in this *ex vivo* condition. However, we view this particular discrepancy as a feature rather than a bug. Because they enable the uncoupling of cell fate from morphogenesis, and of morphogenesis in one tissue from another, our explants are uniquely suited to address the question of tissue-specific C&E. We leveraged this ability to distinguish NE-from mesoderm-specific extension to define a novel role for BMP in C&E of the NE that was not identified through studies of intact embryos alone. Explants similarly enabled identification of distinct temporal signaling dynamics that promote tissue-specific morphogenetic programs.

But why would different times of signaling onset affect each tissue differently? In fact, the temporal dynamics that are critical for morphogenetic outcomes in explants directly reflect signaling events during gastrulation *in vivo*. For example, the high Nodal activity at or before 4 hpf that induces mesoderm extension in explants mirrors *in vivo* Nodal signaling that begins around 4 hpf in the dorsal margin (*50, 53*), the region fated to become (robustly extending) chordamesoderm. Nodal signaling continues within the future mesoderm at the embryonic margin until gastrulation begins (Figure 7A). Once the mesoderm is internalized during gastrulation at 6 hpf and beyond, the chordamesoderm continues to express Nodal ligands as it takes up position beneath the dorsal ectoderm, where it activates Nodal signaling within the overlying NE (*50*) (Figure 7B) that begins to converge and extend shortly thereafter (*3, 87*). Hence, the dynamics of gastrulation create a delay in the reception of Nodal within the NE compared with the mesoderm, both of which later undergo C&E simultaneously (*2, 3*). We speculate that by activating Nodal after 4 hpf (either with opto-Nodal or *ndr2*), explants are denied the early signal that would induce chordamesoderm C&E and receive only the later cue for NE C&E, yielding NE-driven extension. These results are consistent with our previous findings that Nodal signaling contributes to explant C&E even after gastrulation onset, and that this extension is largely NE-driven (*32*).

**Figure 7:**
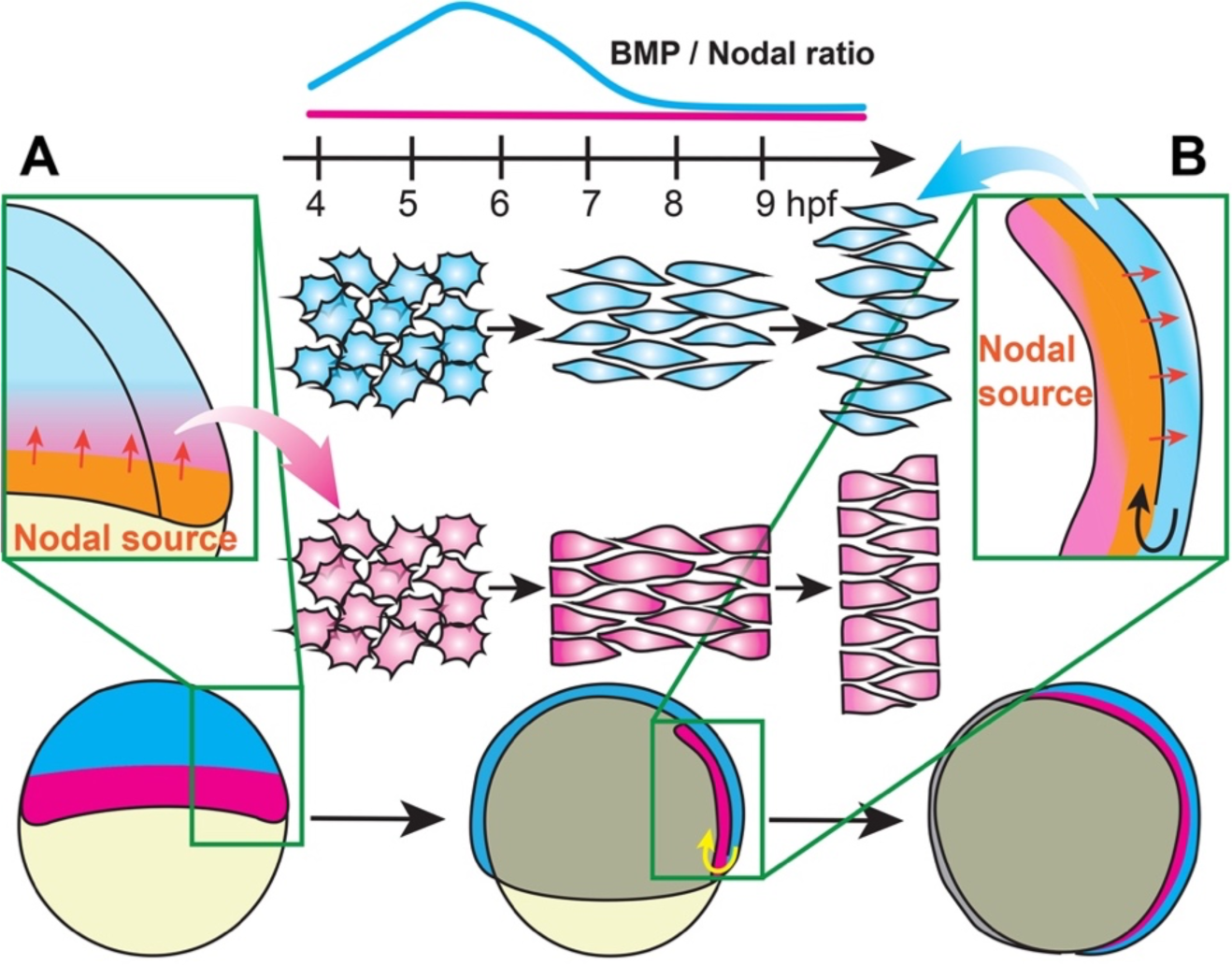
A model for temporal signaling dynamics in tissue-specific gastrulation morphogenesis. During late blastula stages, the embryonic margin is a source of Nodal ligands that activate signaling within the future mesoderm (**A**). During gastrulation, internalized mesoderm continues to express Nodal ligands, serving as a Nodal source that activates signaling within the overlying neuroectoderm (**B**). This creates a delay in the reception of Nodal signals by the NE compared with the mesoderm, resulting in a period of low BMP/Nodal ratios in the mesoderm and high BMP/Nodal ratios in the NE (top) that promote distinct C&E modes in each tissue.

### Cell autonomy of C&E regulation by BMP/Nodal

One notable feature of our explant system is that the NE undergoes C&E even when mesoderm C&E is blocked by BMP activity. This is distinct from previous findings both *in vivo* (*70*) (discussed further below) and in other explant models. It was recently shown, for example, that increased BMP activity reduced C&E of all tissues within whole-blastoderm (pescoid) explants (*29*), similar to intact embryos (*70*) and *Xenopus* explants (*72*). By contrast, increased BMP signaling in our animal pole explants prevented mesoderm extension but enabled NE C&E. Together with our *in vivo* data showing a cell-autonomous increase in ectoderm cell polarization upon BMP activation, this demonstrates a primary and ectoderm-specific role for BMP in promoting C&E cell behaviors. This is in striking contrast to the role BMP plays in the mesoderm, where elevated BMP activity was shown to cell-autonomously inhibit C&E cell behaviors (*70*). The downstream molecular mechanisms by which BMP promotes C&E behaviors in one cell type and inhibits them in another will be important topics for future research.

It was previously reported that when mesoderm C&E was inhibited by high BMP levels, ectoderm cells also exhibited reduced C&E (*70*). Furthermore, the ventral side of the embryo naturally exhibits high BMP signaling and comprises a “no convergence no extension zone” that prevents C&E of all cell types (*70, 88*). These findings raise questions of why, if BMP promotes NE C&E, does the ventral ectoderm not converge and extend in the embryo? One possible explanation is the inter-dependence of mesoderm and NE movements *in vivo*. Increased BMP activity was shown to reduce cadherin-dependent cell adhesion on the ventral side of zebrafish gastrulae (*89*), creating an adhesion gradient that reduces traction of migrating cells ventrally and drives cell displacement dorsally. It is therefore possible that ventral ectoderm cells “attempt” to exhibit C&E behaviors *in vivo* but because they cannot gain traction on their ectodermal or mesodermal neighbors, they are unable to produce the force necessary to drive morphogenesis. How increased BMP activity in ectoderm cells might overcome this barrier is as yet unclear. The reduction of cell adhesion by BMP was also recently shown to regulate migration of the overlying mesoderm by modulating tissue viscosity (*90*), highlighting a potential role for tissue mechanics in NE C&E. Finally, BMP may regulate C&E cell behaviors through feed-forward or feedback signaling, or by cross talk with other signaling pathways, either cell-autonomously or non-autonomously. For example, we cannot rule out a non-autonomous effect of BMP activation in transplanted ectoderm on the host mesoderm that contributes to hyper-extension of embryos. One caveat to our animal pole transplants is that they do not produce embryos in which the entire ectoderm compartment is derived from one condition and the mesoderm compartment from another, meaning that host ectoderm must be considered when interpreting our results as well. Even so, these transplants enable investigation of cell-, and to a certain extent tissue-, autonomy *in vivo* because transplanted animal poles are highly unlikely to contribute to mesoderm. While additional non-autonomous effects are possible, our transplant data indicate that BMP promotes C&E cell behaviors cell-autonomously within the ectoderm.

### Effects of cell fate on C&E

Nodal is crucial for specification and patterning of the mesoderm, with certain mesoderm types exhibiting stronger C&E (chordamesoderm) than others (lateral mesoderm) (*12, 70*). One may hypothesize that the extending vs. non-extending mesoderm phenotype seen in *CA-acvr1b\** and *ndr2* explants, respectively, could result from specification of different types of mesoderm with different propensities for C&E. Our data do not support such a hypothesis, as both explant conditions express markers of the same tissues (Figure 1), suggesting that mesoderm type is not the determining factor of these distinct morphogenic outcomes. However, we cannot rule out subtle difference in cell types that may contribute to the mesoderm’s ability to extend.

This may also be true for tissue-autonomous vs. non-autonomous NE extension. We observed that the graded expression of *sox2* in *CA-acvr1b\** explants differs from the striped expression induced by *ndr2,* which may underlie differences in patterning that could influence NE C&E. For example, NE cells within each explant condition may possess distinct anterior-posterior positional identities, which are both necessary and sufficient for mesoderm C&E (*91*). We show that both *CA-acvr1b\** and *ndr2* explants express the anterior NE marker *otx2*, but additional neural patterning within explants remains to be examined. Such patterning depends on the interaction of multiple signaling pathways: while BMP antagonism induces anterior NE, FGF signaling is required for posterior NE development (*47, 48, 92–94*). Indeed, the combination of BMP antagonism and FGF activity induces extension of *Xenopus* animal cap explants in the absence of mesoderm (*47, 48*), suggesting that posterior positional identities may promote NE C&E. Understanding how neural patterning influences its tissue-autonomous morphogenesis, and how this interfaces with temporal signaling dynamics, remains to be elucidated.

## MATERIALS AND METHODS

### Zebrafish

Adult zebrafish were maintained through established protocols (*95*) in compliance with the Washington University and Baylor College of Medicine Institutional Animal Care and Use Committees. Embryos were obtained through natural mating and staging was based on established morphology (*96*). Studies were conducted using WT and Tg(*79*) (*79*) embryos of AB and AB* strains. Fish were crossed from their home tank at random and embryos were chosen for injection and inclusion in experiments at random.

### Microinjections of synthetic mRNA and guide RNAs

Single-celled embryos were placed in agarose molds (Adaptive Science tools I-34) and injected with 0.5-2 nL volumes using pulled glass needles (Fisher Sci #50-821-984). Doses of RNA per embryo are as follows: 0.5 pg CA-*acvr1b*\* (*80*), 10 and 100 pg *ndr2* (*97*), 10.4 pg Alk3/10.4 pg Alk8/17.8 pg BMPR2A (*61*), 5 pg acrv1b-Cry2/5 pg acvr2b-CibN (McNamara et al, in preparation), 50 pg CA-*BMPR1A* (*85*), 100 pg *admp* (*86*), and 50 pg *chrd* (*98*), 200 pg *membrane-cherry*. All mRNAs were transcribed using the SP6 mMessage mMachine kit (Fisher Sci #AM1340) and purified using Biorad Microbiospin columns (Biorad #7326250). Two guide RNAs targeting *chordin* and Three guide RNAs targeting *tyrosinase* were transcribed using T7 RNA polymerase (NEB #M0251s) from DNA templates made from the following primers for *chordin*: (1:gaaattaatacgactcactataGGTGGCCGCTTTTACTCTCgttttagagctagaaatagc, 2:gaaattaatacgactcactataGGGACCGGATCGTCGCAGGTgttttagagctagaaatagc) or *tyrosinase* (1: gaaattaatacgactcactataGGTGCGCCGGAAACTACATGgttttagagctagaaatagc, 2: gaaattaatacgactcactataGGACTGGAGGACTTCTGGGGgttttagagctagaaatagc, 3: gaaattaatacgactcactataGGGTGTACGATTTATTCGTGgttttagagctagaaatagc) (as described in (*99*)). A cocktail of guide RNAs for each target (1 μL each) was pre-incubated with Cas9 protein (NEB #M0646M) and 0.68uL KCl (as described in (*100*)). 1 nL of each complex was injected into embryos at the single-cell stage.

### Immunofluorescent staining

Embryos and explants were stained for phosphorylated Smad2 (Cell Signaling #18338S) and phosphorylated Smad5 (Cell Signaling #13820S) as described in Van Boxtel et al., 2015. Briefly, samples were fixed overnight in 4% paraformaldehyde (PFA) (VWR #101176-014) in phosphate-buffered saline (PBS), Rinsed in PBS (Thermo/Fisher #14190250) + 0.1% Tween-20 (Thermo-Fisher #23336-2500) (PBT), dehydrated into methanol, and stored at -20C overnight. Samples were rehydrated into PBS + 1% Triton X-100 (PBTr) (VWR #97063-864) and incubated with ice-cold acetone at -20C for 20 minutes. Samples were blocked in PBTr + 10% newborn calf serum (Thermo/Fisher #26010066) for one hour and then incubated overnight at 4C with anti-pSmad2/3 or anti-pSmad1/5 at 1:1000 in block. Samples were rinsed with PBTr and incubated with Alexa Fluor 488 anti-Rabbit IgG (Thermo/Fisher #PIA32731) at 1:1000 and 4′,6-diamidino-2-phenylindole, dihydrochloride (DAPI) at 1:5000 overnight on a 4C shaker. Embryos and explants were rinsed in PBTr prior to mounting in 0.5% low-melt agarose (Thermo/Fisher #16520100) for confocal imaging.

### Whole mount in situ hybridization

Antisense riboprobes were transcribed using NEB T7 or T3 RNA polymerase (NEB #M0251s and Fisher Science #501047499) and labeled with digoxygenin (DIG) NTPs (Sigma/Millipore #11277073910) or fluorescein NTPs (Sigma/Millipore #11685619910). Whole-mount in situ hybridization (WISH) was performed according to (*101*) with modifications. Embryos and explants hybridized with anti-fluorescein probes were washed extensively in MABT (100 mM maleic acid, 150 mM NaCl, 0.1% Tween-20, pH 7.5) and placed in RBR buffer (20% Goat serum, 2% Roche Blocking Reagent in 1x MAB) on the orbital rocker for 1 hour. RBR buffer was removed and replaced with 1:5000 anti-fluorscein-AP antibody (Roche #11681460001) in RBR buffer and incubated on the 4C orbital shaker overnight. The following day, samples were washed extensively in MABT buffer and staining buffer (PBT, 100 mM Tris [pH 9.5], 50 mM MgCl2, 100 mM NaCl) was added. Staining Buffer was removed from samples and developing solution was added until the desired strength of staining was reached. Samples were placed in stop buffer (10 mM EDTA in PBT) before imaging.

### Blastoderm explants

Blastoderm explantation was performed according to (*33*). Embryos were dechorionated with pronase (1 mL of 20 mg/mL stock in 15 mL 3X Danieau’s) at 128 cell stage and washed with egg water and 0.3X Danieau’s. Explants were cut using Dumont #55 forceps (Fisher Science #NC9791564) on an agarose coated 60 mm X 15 mm plate filled with 3X Danieau’s. After cutting, explants were allowed to heal in the 3X Danieau’s for 5 minutes before being placed in agarose coated 6-well plates filled with explant media (DMEM/F12 Thermo #11330032, 3% Newborn calf serum, 1:200 penicillin-streptomycin). Explants were collected and fixed in 4% PFA when stage matched intact embryos reached 4-somite stage (∼12 hpf).

### Animal pole transplantation

Animal pole transplants were performed similarly to blastoderm explants. Dechorionated embryos were placed in an agarose mold (Adaptive science tools #PT-1) containing 3X Danieau’s. The top half of the donor embryo was cut and placed next to the receiver embryo. The receiver embryo was cut in a similar manner. The cut side of the donor explant was pressed onto the cut side of the receiver embryo and held for a few seconds until they were securely together. The newly transplanted embryos were then placed into a well from the mold and allowed to heal for 15 minutes before transferring into an agarose coated 6-well plate or an agarose coated 35mmx10mm dish filled with 0.3X Danieau’s solution.

### Optogenetic Tools

All embryos injected with mRNAs encoding optogenetic receptors were placed in the dark no later than 4 cell stage and samples were manipulated and mounted under red lamps to avoid activation prior to the experiment’s onset. Samples that were kept as dark controls were placed in a foil-wrapped box within the incubator to prevent activation. Embryos or explants were exposed to blue illumination in a 6-well plate with 1mL of agarose and 4mL of 0.3X Danieau’s or explant media respectively. 6-well plates were placed 10 cm under blue LED light strips (Amazon, L1012V-MC1-0330-ECLBK-K-N) in a 28.5C bench-top incubator. For opto-BMP explants, 6-well plates were placed directly (<1cm) under blue LED light strips.

### RNA Sequencing

RNA for sequencing was isolated from 40 pooled explants of each condition (*ndr2*, *CA-acvr1b\**, and uninjected) at each of seven stages (sphere, 30% epiboly, 50% epiboly, shield, 75% epiboly, 90% epiboly, and 2 somite) from each of three independent experiments. Total RNA was isolated using Trizol reagent (Thermo Fisher #15596018), then purified by sodium acetate precipitation and eluted in 10 mM Tris pH7.5. Samples were submitted to Novogene (Sacramento, CA) for library preparation, including polyA enrichment. Libraries were sequenced using an Illumina NovaSeq 6000 to obtain paired-end 150bp reads. Data quality was assessed using FastQC. Low quality basepairs and adapters were trimmed using trimGalore. Data were mapped onto the zebrafish danRer11 genome assembly using STAR (*102*). Gene expression was assessed using featureCounts (*103*). Differentially expressed transcripts were determined using the R packages EdgeR (*104*) and RUVr (*105*), with significance achieved for FDR<0.05 and fold change exceeding 1.5x.

### Microscopy

Immuno-stained embryos and explants were mounted in 0.5% low-melt agarose (Thermo/Fisher #16520100) in glass bottomed 35 mm petri dishes (Fisher Sci #FB0875711YZ) for imaging using a Nikon ECLIPSE Ti2 confocal microscope equipped with a Yokogawa W1 spinning disk unit, PFS4 camera, and 405/488/561nm lasers (emission filters: 455/50, 525/36, 605/52). For explants, 60 μm z-stacks were obtained with a 2 μm step using a Plan Apo Lambda 20X lens. For opto-Nodal embryos, 72 μm z-stacks were obtained with a 2 μm step using a Plan Apo Lambda 10x lens. For cell shape of Opto-BMP embryos, 8-29 μm zstacks were obtained with 2 μm step using a Plan Apo Lambda 40X lens. Images of WISH-stained embryos and explants were taken with a Nikon Fi3 color camera on a Nikon SMZ745T stereoscope.

### Image analysis

ImageJ/FIJI was used to visualize and measure all microscopy data sets. Researchers were blinded to the conditions of all image data using the *blind_renamer* Perl script prior to analysis.

### Morphometric Measurements

To measure the length/width ratios of explants, we divided the length of the segmented line along the midline of the explant (accounting for curvature) by the length of the perpendicular line spanning the width of the explant near its midpoint. This measurement technique was also used for mesoderm domains where the length of the segmented line along the midline of the *tbxta* stain was divided by the length along the midline along the whole explant. To measure the length of whole embryo embryonic axis, lateral-view images were collected of each embryo, and a segmented line was drawn along the embryo curvature from the anterior *dlx3b* expression domain to the posterior *tbxta* expression domain. The distance between the prechordal plate and the notochord was measured from dorsal-view images of each embryo. A line was drawn from the top of the *tbxta* expression domain to the bottom of the *hgg1* expression domain. All embryos were stage matched with *egr2b* expression.

### Individual Cell Measurements

Within live embryos expressing membrane Cherry, a projection of several z-planes through the ectoderm was chosen for each condition when embryos were at bud stage. To measure cell orientation and elongation, all samples were placed in their respective tissue of interest to be analyzed and then blinded. A fit eliipse was used to measure the aspect ratio and orientation of the cell’s major axis.

### pSmad2/3 and pSmad1/5/8 immunostaining quantification

DAPI z-stacks were converted into 3-dimensional masks, which were subtracted from pSmad z-stacks to create 3D stacks of only nuclear pSmad labeling. FIJI’s ‘3D Objects counter’ plugin was then used to detect the location and pSmad fluorescence intensity of all nuclei in a given explant. All nuclei with fluorescence intensities above a threshold background level were categorized as pSmad-positive. We then graphed the maximum pSmad fluorescence intensity among pSmad-positive nuclei.

### Statistical Analysis

Graphpad Prism 10 software was used to perform statistical analyses and generate graphs for all data analyzed. Datasets were tested for normality prior to analysis and statistical tests were chosen accordingly. The statistical tests used for each data set are noted in figure legends.

## ACKNOWLEDGEMENTS

We thank Dr. Lila Solnica-Krezel for generously sharing plasmids, fish lines, support, and wisdom. Animal care was provided by the BCM Center for Comparative Medicine and Washington University Zebrafish Facility. We thank Dr. Katherine Rogers for generously providing opto-BMP receptors. Also, Drs. Ann Sutherland, Ray Keller, Dave Shook, and John Wallingford for helpful discussions and Williams’ lab members for technical assistance, support, and feedback on this project.

## DATA AVAILABILITY

Raw RNA-sequencing data are available on NCBI GEO accession number GSE246158 and will be made public at the time of publication.

